# Reading out olfactory receptors: feedforward circuits detect odors in mixtures without demixing

**DOI:** 10.1101/054247

**Authors:** Alexander Mathis, Dan Rokni, Vikrant Kapoor, Matthias Bethge, Venkatesh N. Murthy

## Abstract

The olfactory system, like other sensory systems, can detect specific stimuli of interest amidst complex, varying backgrounds. To gain insight into the neural mechanisms underlying this ability, we imaged responses of mouse olfactory bulb glomeruli to single odors and hundreds of mixtures. We used this data to build a model of mixture responses that incorporated nonlinear interactions and trial-to-trial variability and explored potential decoding mechanisms that can mimic mouse performance from our previous study (Rokni et al., 2014) when given glomerular responses as input. We find that a linear decoder with sparse weights could match mouse performance using just a small subset of the glomeruli (~ 15). However, when such a decoder is trained only with single odors, it generalizes poorly to mixture stimuli due to nonlinear mixture responses. We show that mice similarly fail to generalize, although they could in principle fully decompose the mixtures. This suggests that mice learn this segregation task discriminatively by adjusting task-specific decision boundaries without taking advantage of a demixed representation of odors.

## INTRODUCTION

In natural environments, the olfactory system must be able to detect behaviorally relevant odors against varying background smells. This task resembles the cocktail party problem in audition where the signals of different sound sources also arrive as a linear mixture at the sensory organ. Similar to our remarkable ability of disentangling sounds sources, mice have been shown to excel at the olfactory equivalent (Rokni et al., 2014). Yet, with increasing number of mixture components this task can get difficult due to the overlapping, combinatorial re-presentation of odorants by receptor neurons (Duchamp-Viret et al., 1999, Koulakov et al., 2007, Rospars et al., 2008, Shen et al., 2013) and sparse, distributed representations in higher olfactory areas (Stettler and Axel, 2009, Wilson and Sullivan, 2011). These representations have been described as synthetic, in the sense that combinations of odorants are thought to be encoded rather than individual components (Gottfried, 2010, Wilson and Sullivan, 2011).

In the main olfactory bulb (OB) of the mouse each sensory neuron expresses only 1 of ~ 1000 types of olfactory receptor proteins (Godfrey et al., 2004, Young and Trask, 2002, Zhang and Firestein, 2002). Each odor will activate a subset of these receptor types and will therefore be identified by this subset; and each receptor type may be activated by many odors, leading to a potentially large overlap in the representation of different odorants (Duchamp-Viret et al., 1999, Malnic et al., 1999, Rubin and Katz, 1999, Shen et al., 2013). Furthermore, mixtures may be represented as nonlinear functions of their parts (Rospars et al., 2008). Because of these factors, it is not clear how the difficulty of detecting individual components relates to the number of background odors. In a previous study, we found that mice can be trained to perform a target-odor detection task and can do so remarkably well even in the presence of many background odors (Rokni et al., 2014). These behavioral results led to the question of how mice solve this task given the nonlinear, noisy and high dimensional input that the glomeruli provide.

Various attempts have been made previously to provide theoretical understanding of how odors can be detected against a background. Several algorithms have relied on differences in timing of arrival of odorants at the sensors (Hendin et al., 1998, Hopfield, 1991, Li and Hertz, 2000). Others proposed that the temporal structure of OB activity provides a concentration and background invariant code for odor identity (Brody and Hopfield, 2003, Galán et al., 2006, Hiratani and Fukai, 2015). Detection of a specific component can also be achieved using Bayesian inference if one assumes prior knowledge of the receptor activation levels for ′all odors′. Such prior knowledge could be implemented in the brain as a ′generative model′, which could be formed by unsupervised learning (Friston, 2010, Hinton, 2007a, Hinton, 2007b). Unsupervised learning could yield ′demixed′, atomic representations of odors in higher olfactory areas - akin to Barlow′s idea that a lot of knowledge about the world is imprinted into the representations of sensory systems (Barlow, 1997). Based on a linear encoding model, Bayesian inference has been shown to work well in conditions where the typical mixture contains few components (Grabska-Barwinska et al., 2013). More generally, many algorithms for blind source separation and inference could be applied to such a generative model (Bell and Sejnowski, 1995, Comon and Jutten, 2010, Haykin and Chen, 2005, Otazu and Leibold, 2011, Tootoonian and Lengyel, 2014). Alternatively, supervised methods, like linear classifiers could be used for discriminative learning (Galan et al., 2006, Bishop, 2006, Berens et al., 2012, Shen et al., 2013). None of these studies, however, attempted to identify the decoding mechanism that mimics mouse psychophysical performance - this is the focus of the current work.

In this study, we sought to find the simplest decoding mechanism that uses experimentally constrained neuronal input and match the ability of mice to detect target odorants (Rokni et al., 2014). We measured representations of single odors and hundreds of mixtures by olfactory receptors to derive a quantitative model of mixture responses, which also included saturation and experimentally-measured trial-to-trial variability. Using this model to generate glomerular responses to arbitrary mixtures, we tested whether simple biologically-plausible read out mechanisms can solve the behavioral task. We find that the simplest multi-glomerular approach, which sums up the weighted activity of different glomeruli and compares this sum to a threshold (a linear classifier), is sufficient to mimic the behavior of mice. Several glomeruli have to be combined, which argues against a strict labeled line encoding model. The quality of the mixture samples used in training the model was important, since a model trained only on single odors only poorly generalizes to mixtures. We tested this prediction in mice and find a similar behavior.

## RESULTS

In the behavioral task, mice were required to report the presence of one of two target odors in random mixtures containing up to 14 odors (Fig. 1A). After learning, mice achieved accuracies close to 100% for few distractors, and performance declined with increasing number of background odors (Rokni et al., 2014). This is a seemingly remarkable feat given that a) mice had to generalize from around 1,000 training trials to mixtures that they had never smelled before (more than 60% are novel in test phase, ~ 50,000 possible mixtures), b) glomerular patterns are highly overlapping, c) mixture responses arise from nonlinear interactions of single odors, and d) odor responses are highly variable from trial-to-trial. To assess how challenging this task actually is, we started by quantifying all the parameters of the input at the level of receptors and then probed the ability of decoders to use glomerular activity patterns to extract the presence of particular target odors.

### Estimating an encoding model for mixtures using empirical data

In earlier experiments (Rokni et al., 2014), we had measured the average glomerular responses to the 16 individual odors in anesthetized mice expressing the Ca^++^ indicator GCaMP3 in all olfactory receptor neurons. The measured signals arise from the collection of sensory axons converging on glomeruli. The number of glomeruli that were responsive to at least one odor in each OB ranged between 63 and 76.

Due to the large number of possible odor mixtures, measuring the neural responses to the whole stimulus set is intractable (> 10,000 mixtures). Therefore, we estimated a statistical model that describes the glomerular responses given a particular odor mixture. We imaged responses to all single components as well as a sample of ~ 500 mixtures in 5 additional OMP-GCaMP3 mice (Isogai et al., 2011, Fig. 1C-L). To estimate the variability of responses each single odorant was presented several times per experiment (7.7 ± 1.7, mean ± SD, Fig. 1F). Over all odor-glomerulus pairs, the average coefficient of variation (CV) was 0.37±0.07 (mean ± SD). However, much of the variability was correlated across glomeruli, probably reflecting causes that are either irrelevant to behaving mice (e.g. anesthesia level and breathing parameters; Blauvelt et al., 2013) or could be easily factored out by neural operations such as normalization. This correlated variability is evident when the responses of all glomeruli to each trial of a specific odor were plotted against the mean response to that odor across trials (Fig. 1G). To estimate the non-correlated variability, we first subtracted the correlated variability from all responses. This was achieved by subtracting from each response pattern its best linear fit to the mean response pattern of the same odor (solid lines in Fig. 1G, see Eqn. (4) in Methods). The timescale of changes in the linear fit parameters was on the order of 10 minutes, suggesting factors such as changes in anesthesia (Fig. 6). For each glomerulus *i* and odor *j*, we then calculated the uncorrelated component of the standard deviation (Eqn. (4)) across trials *t*, and plotted it against its mean response amplitude *O_ij_*. The non-correlated CV for each experiment was calculated as the slope of the linear relationship between the non-correlated SD and response amplitude (Fig. 1H). Non-correlated CVs ranged between 0.08 and 0.13 (0.099±0.019, mean ± SD, n=5 mice). This value is higher than expected based on the CVs measured in single rat receptor neurons in vivo (Duchamp-Viret et al., 2005) indicating that our measure of variability is probably conservative.

To estimate how glomerular responses to mixtures relate to their responses to mixture components, we imaged a sample of ~ 100 mixtures per experiment (98±39, mean ± SD, n=5) and compared the responses with the sum of the responses to individual components of the mixture (Fig. 1I-L). Most mixture responses could be estimated rather well by the linear sum of individual component responses (Fig. 1I). However, beyond a certain value, mixture responses saturated (Fig. 1J-L). Saturation in odor responses may be due to several factors such as odor-receptor protein interaction, receptor neu-ron firing, and saturation of the Ca^++^ indicator. We could not disambiguate which of these contribute to saturation in our data set, and since receptor ligand interactions are well known to be saturating with increasing ligand concentration (Firestein et al., 1993, Grosmaitre et al., 2009, Reisert and Matthews, 2001), we conservatively assume that saturation is a real characteristic of olfactory receptor neuron′s odor responses and use a saturating curve in our model for mixture responses.

**FIG. 1.**
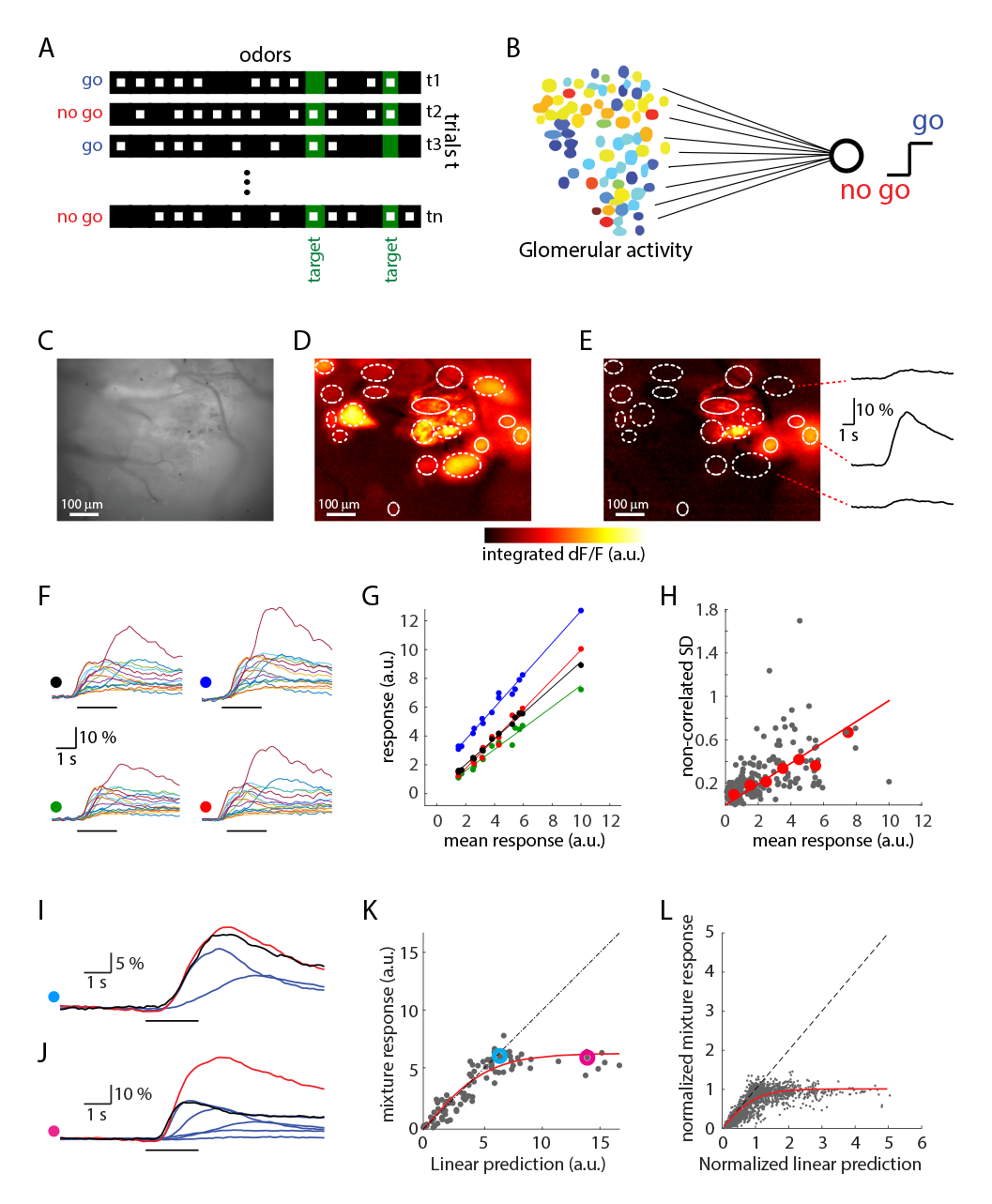
Variability and summation of glomerular odor responses. (A) Task structure. In each trial an odor mixture is randomly generated from a pool of 16 odors. The task is to categorize based on the presence of one of two targets (green). Filled squares denote present odors and empty squares denote absent odors. (B) The classifier reads the glomerular responses as inputs and attempts to report the presence of the targets. (C) Raw, temporal average fluorescence in one imaged area. (D) Superposition of the *dF/F* responses to the 16 odors for same area as in C. Ellipses show putative glomeruli. (E) Average response to ethyl propionate. Time course of fluorescence *dF/F* changes shown for three glomeruli at right. (F) Responses to four presentations of methyl tiglate. Each panel shows one trial for all glomeruli. (G) These four trial-responses to methyl tiglate are plotted against the mean response. Colors represent trials as indicated by colored dots in F. Lines are linear fits. (H) The trial-by-trial standard deviation after removal of correlated variability (see Methods) is plotted against the mean response (gray dots) for each glomerulus-odor pair. Red dots are the median of mean-response-binned data. Red line is a linear fit to the data (forced through origin). (I) An example of linear summation of mixture components. Shown are the average responses to individual components (blue), the response to the mixture (black), and the linear sum of the responses to the individual components (red). (J) An example of sub-linear summation of mixture components. Colors as in I. (K) All mixture responses are plotted against the linear sum of the responses to individual components for one single glomerulus. Colored data points show the examples in I and J. Red line depicts the best sigmoid function fit (Eqn. (5), see Methods). (L) Same as I with data pooled across glomeruli and mice. Both the linear sum and the experimental data were normalized for each glomerulus to the saturation value of the fitted sigmoid function.

These mixture responses can thus be summarized by the following encoding model that describes the activity vector *R* of glomerular responses in trial *t* (see Methods):

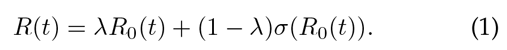

Thereby *λ* ∈ [0,1] interpolates between the fully linear model *R*_0_(*t*), and *σ* denotes the saturating nonlinearity giving rise to the saturated model *σ*(*R*_0_(*t*)) (Eqn. (5)). The linear encoding part is given by

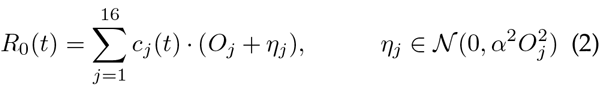

where *O*_*j*_ is the glomerular response pattern for the *j*th odor, *c*_*j*_(*t*) a binary variable denoting the presence of odor *j* in trial *t*, and the noise term *η*_*j*_ is normally distributed with a mean of zero and a variance of 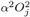. The factor *α* controls the coefficient of variation. Our empirical estimate for this parameter is ~ 0.1. We will refer to the sequence of *c*_*j*_(*t*) during an experiment as the trial structure.

In addition to introducing nonlinear mixing, parametrized by *λ*, we also systematically varied levels of trial-to-trial variability *α* to study its effect on decoding performance.

### Performance of optimal linear decoder for measured mixture model

The simplest decoder at the population level is a linear readout, similar to a single perceptron (Rosenblatt, 1958). Such a linear readout weighs the activity of individual glomeruli *R*_*i*_(*t*) by synaptic weights *w*_*i*_ and predicts the presence or absence of a target when the summed product is larger or smaller than a threshold *θ*, respectively. Mathematically, the output is given by

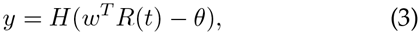

where *y* is the binary output, *H* the Heaviside function, which is 1 for positive arguments and 0 otherwise and *T* denotes the transpose of the weight vector *w*.

In our published work (Rokni et al., 2014), there were thirteen mice and 1, 500 - 3, 500 trials per mouse (2, 308 ± 625, mean ± SD) with uniform distractor distribution, i.e. the number of odors per trial was equally distributed between 1 and 14. For each mouse this segregation task can be summarized by the trial statistics *C*_*j*_(*t*), which is a binary variable denoting the presence of odor *j* in trial t and the ideal behavior *r*(*t*). For each animal´s trial statistics, we calculated the optimal linear weights *w*_*OLE*_ that minimize the mean squared deviations between the ideal behavior *r*(*t*) and the output of the linear decoder *w*^*T*^*R*(*t*) (see Eqn. (9)) and the optimal decision threshold and the optimal decision threshold *θ* that maximizes the performance these weights (*w*_*OLE*_), on a random training set of 80% of the trials with glomerular patterns drawn from the encoding model (Eqn. (1)). This decoder, which we call the OLE (for optimal linear estimator) was then tested on the remaining 20% of trials (Methods). We repeated this procedure 20 times and report the average performance on the test sets.

**FIG. 2.**
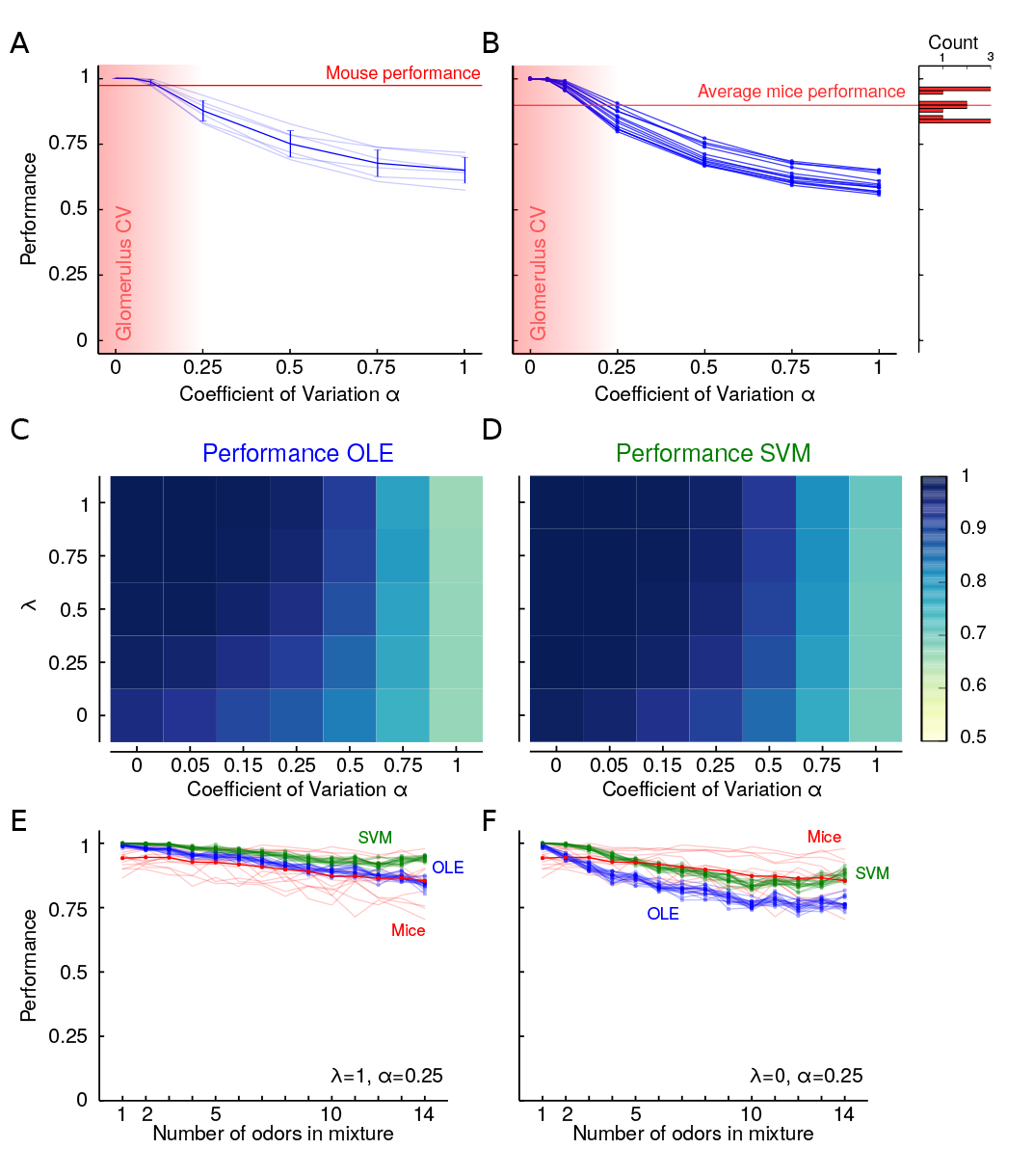
Decoding performance of linear readout and SVM for mixtures with trial-to-trial variability and nonlinear mixtures. (A) Performance of optimal linear readout in detecting one target pair (isobutyl propionate and allyl butyrate) is plotted as a function of trial-to-trial variability (*α*). The curves show the performance of one decoder based on each of the six measured glomerular maps and their average (± SD). The mouse performance for these targets is shown as horizontal line. The estimated range of the glomerular noise is highlighted by the shaded, red region (*λ* = 1). (B) Average decoding performance for all 13 pairs of target odors. The average performance of all mice is indicated by a horizontal line and the distribution of experimental performances is shown to the right (*λ* = 1). (C) Average performance for data from all mice of optimal linear decoder for varying noise levels and nonlinearities from *λ* = 1 (linear case) to *λ* = 0 (experimentally measured saturation). (D) Average performance for data from all mice, using SVM with radial basis function for mixture data as in panel C. (E) Performance of OLE and SVM as a function of the number of odors in the mixture. Performance curves for 13 mice and their average in red. The average performance curves for the SVMs and OLEs are shown for the same 13 target odor pairs for linear mixture model (*λ* = 1) with *α* = 0.25. Those are the conditions for which the experimental and model performance curves are best correlated (Figures S4C-D). (F) Same as (E) but for mixtures with saturation (*λ* = 0).

Performance was perfect without trial-to-trial variability, *α* = 0 (Fig. 2B, *λ* = 1). Increasing *α* gradually decreased performance, which reached ~ 65% for *α* = 1 (Fig. 2A and B). Similar trends were obtained using the glomerular patterns from different imaging datasets (Fig. 2A) and for different target odors (Fig. 2B). For a noise level similar to our experimental estimate (~ 0.1), most linear readouts performed better than corresponding mice detecting the same targets (Fig. 2B). Thus, considering only the trial-to-trial variability and mixing of odors without saturation, a simple linear decoder is sufficient to match or exceed mouse performance, although it had access only to the time-averaged response of a limited set of dorsal glomeruli. Since the specific choice of data had little effect on classifier performance, we focused on a particular imaging set with 72 glomeruli.

Next we introduced glomerular saturation by varying the parameter *λ* in Eqn. (1). As expected, the average performance of the optimal linear decoder for all target odors declined with both increasing nonlinearity and noise (average performance for all target pairs is shown in Fig. 2C, an example target pair is shown in Fig. 7A). When noise levels are low (*α* < 0.1) and responses are not fully saturating (*λ* > 0), the mean decoder performance is above 98%. When considering saturating glomeruli (*λ*=0) and the experimental estimate of noise (*α*=0.1), performance drops to 91.6%, which is comparable to the average mouse performance of 90.3%. This demonstrates that the ′cocktail party problem′ is almost linearly separable at the level of glomeruli.

### Performance of support vector machine (SVM) for noisy, nonlinear mixtures

Since the glomerular activity patterns, even in the linear noisy mixture case, *R*(*t*) are given by a superposition of multiple Gaussian random vectors, the optimal decision boundary might be highly nonlinear and not merely a single hyperplane. When the odor patterns are nonlinearly mixed, the geometry of the decision boundaries becomes even more intricate. A decoder that is able to flexibly adjust its decision boundaries without constraining them to be a single hyperplane may therefore perform considerably better than the simple linear readout. One powerful class of decoders with such properties are SVMs with radial basis functions as kernel (see Methods). The decision boundaries for such SVMs are formed by sums of Gaussians and can therefore in the limit approximate almost any decision boundary (Muller et al., 2001, Vapnik, 1999).

We trained SVM decoders with the same crossvalidation procedure as described for the optimal linear decoder (see Methods). As expected from their versatility, SVMs were more accurate than the linear readout. Fig. 2D. shows the performance of the SVM for the same target odors as the optimal linear decoder in Fig. 2C. When noise levels are low (*α* < 0.1) and responses are not fully saturating (*λ* > 0), the SVM performance was above 99.5%. When considering saturating glomeruli (*λ* = 0) and the experimental estimate of noise (*α* = 0.1) performance drops to 94.7%, outperforming mice and making about 40% fewer mistakes than the optimal linear decoder. Similarly, for other conditions, SVMs make fewer errors than the optimal linear decoder and have substantially better performance for low noise levels irrespective of *λ* (Fig. 2C-D).

Given that the overall performance of both optimal linear decoder and SVM broadly resembled that of mice, we investigated whether they make similar errors. Specifically we asked whether the dependence of performance on the number of mixture components is similar to the monotonic decline seen in mice (Fig. 2E and F). We refer to this relationship as the performance curve. We calculated the average correlation coefficient (*r*) between the performance curves of the classifiers for specific odor targets and their corresponding mice. The average correlation was maximal for the fully saturated mixture model (*λ*=0) with a noise level of *α* = 0.25 for both the optimal linear decoder (*r* = 0.42 ± 0.01, mean ± s.e.m., *n* = 13 mice) and SVM (*r* = 0.40 ± 0.01, mean ± s.e.m., *n* = 13 mice; Fig. 2C and D). When scrutinizing the shape of the performance curves, it is important to note that the number of possible mixtures varies with the number of components in the mixture. For instance, there are many more possible mixtures of 7 components than there are of 1 or 14 components. Therefore, specific stimuli on the edges of the performance curve repeat much more, offering more training samples for the classifier. For this reason, one may expect that classifier (and mouse) performance curves will not be monotonic, but could have a minimum at the midrange of mixtures. We find, however, that despite the different computational capabilities of SVMs and the OLE, performance for both decoders decreases as the number of components in a mixture increase, for realistic estimates of noise and nonlinearity. This suggests that the overlapping and nonlinear mixing of noisy glomerular patterns, rather than limitations of the readout mechanism, is a fundamental bottleneck for this task.

Taken together, although the linear decoder is not the optimal decoder, its performance is comparable to mice for our conservative estimates of noise and saturation levels. We therefore used this linear decoder, rather than the SVM, for further analysis. Nevertheless, it is conceivable that the many parallel pathways from the OB could implement nonlinear decoding schemes such as the SVM; however, for this task nonlinear decision boundaries are not necessary.

### Performance of sparse, linear decoder for noisy, nonlinear mixtures

The linear decoders considered so far had no restrictions on the number of glomeruli used. Inspired by the observation that the projections from the OB to the piriform cortex are sparse and disperse (Ghosh et al., 2011, Miyamichi et al., 2011, Sosulski et al., 2011), we wondered how well sparse, linear readouts would perform the task. We turned to logistic regression, which is an efficient, binary classification algorithm that can be readily interpreted from a neuronal point of view as a linear readout neuron with nonlinear firing (Berens et al., 2012, Bishop, 2006). Similar to the OLE, the optimal logistic regression (OLR) is found by minimizing the number of misclassifications on the training set. Additionally, we employed a regularization term that punishes non-zero readout weights by adding the sum of all absolute values of the weights to the number of misclassifications (L1 - regularization, Eqn.(12)). By weighing this regularization term with a constant 1/*C* that can be systematically varied, one can bias the OLR to exhibit varying degrees of sparseness. This regularization can be thought of as an additional cost term for neuronal wiring, and that the OLR attempts to maximize performance while minimizing wiring.

When the cost for wiring dominates, the performance of the OLR drops to chance level and at the other extreme performance asymptotes to levels similar to the optimal linear. Between these two extremes performance strongly depends on the number of nonzero readout weights (Fig. 3A). However, the overall effect of readout sparseness on performance was mild; this is not a simple consequence of the glomerular activity itself being sparse, since most glomeruli are activated by at least one odor. For our estimated noise level of 0.1, only 10-20 glomeruli are sufficient to match the performance of mice. To reach 90% of the asymptotic performance only 15.7 ± 0.9, mean ± s.e.m. (*n* = 13 mice) were necessary (Fig. 3A). The corresponding readout weights for 90% of the asymptotic performance and the asymptotic performance are shown in Fig. 3B. The OLR is trained merely based on the sampled mixtures vectors *R*(*t*), and the ideal response for the fraction of training trials, but has no explicit knowledge of the identity of the target odor or the glomerular activity patterns of the individual components. As indicated above, there are at least an order of magnitude more mixture stimuli than stimuli that a given mouse was exposed to. Despite these differences in trial statistics, classifiers trained on trial statistics from different mice with the same target odors converged to similar weights in both the sparse and the dense case (Fig. 3B - see odor pairs F, G & H, for example). When mice had different targets, the readout weights were also distinct (Fig. 3B). These weights are not simply a reflection of the glomerular patterns of the target odors, but also take into account glomerular patterns of distractors (Fig. 8A). Conversely, attempting to use the target-odor patterns as readout weights results in poor performance - even for just one target odor such a template matching algorithm is about 20% worse (Fig. 8B-C).

**Fig. 3.**
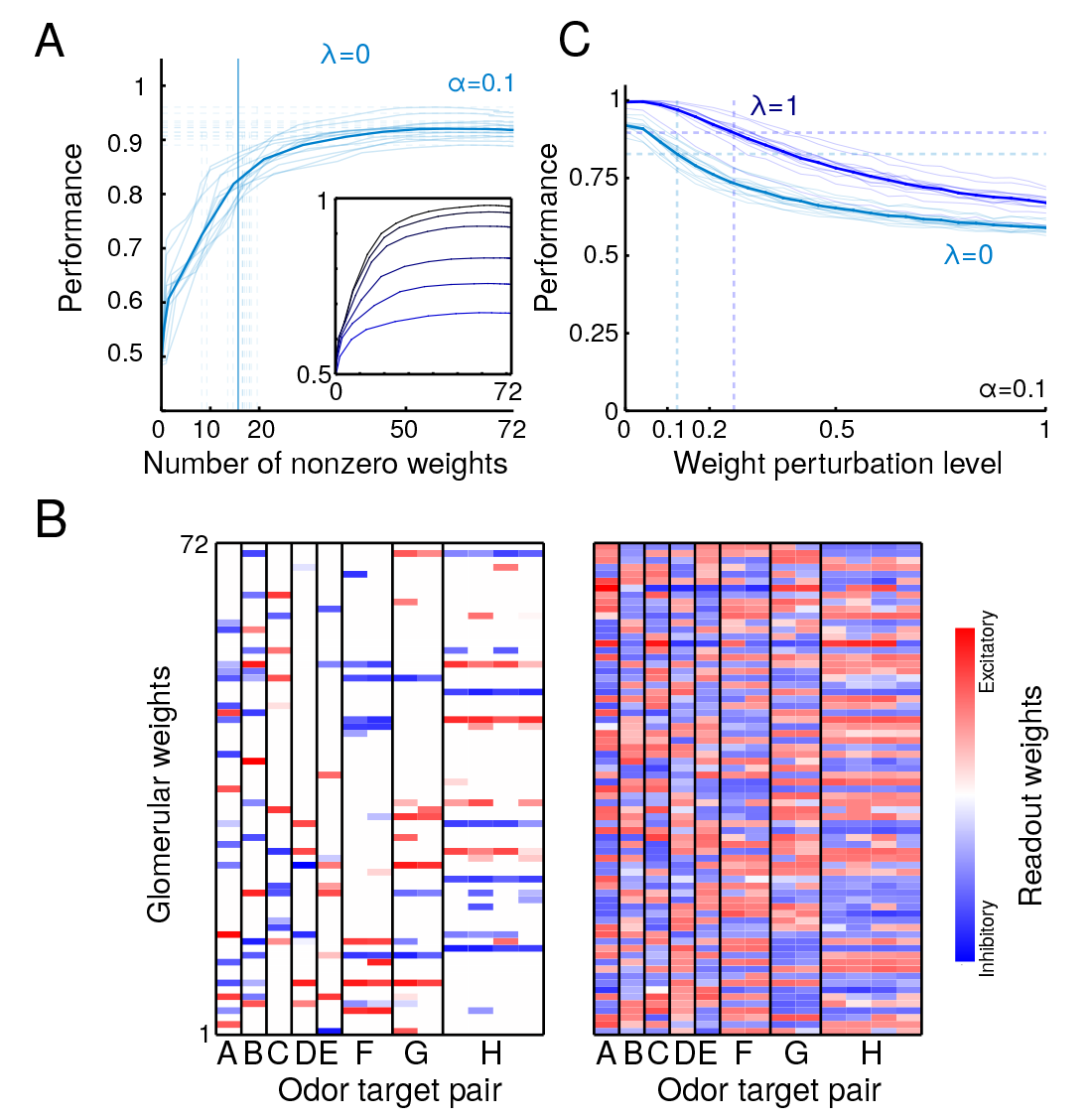
Performance of sparse linear readout (OLR) (A) Performance versus number of non-zero weights of the OLR (*λ* = 0). Each faded line is for data from one animal and solid line depicts the average. The horizontal, dashed lines indicate asymptotic performance and the vertical dashed lines indicate the corresponding minimal number of glomeruli to achieve 90% of that performance per animal. Their average is highlighted as solid vertical line. Inset: Average performance vs. number of non-zero weights of the OLR for varying noise levels (*α* = 0, 0.05, 0.1, 0.25, 0.5, 0.75, 1, from top to bottom). (B) Example readout weights obtained for 90% of asymptotic performance (left) and asymptotic performance (right). Several of these 13 mice had the same target odor pairs, as indicated by grouping under the same letter. OLR learns similar readout weights from different trial statistics for the same target odors. Across different target odors, the readout weights vary substantially - but the weights are similar for the same target odors. (C) Performance of OLR versus perturbation level of weights for linear and nonlinear mixing with *α* = 0. 1. Each thin line is the average for 20 random perturbations per mouse, and the thick line is the average across mice. The horizontal lines indicate 90% of the unperturbed weight performance for either condition (*λ* = 0,1). The vertical lines indicate the perturbation level where the performance drops below 90% of the unperturbed weight performance.

The saturation also affects which and how many glomeruli are read out by the OLR (Fig.9A-B). For linear mixing (*λ* = 1), only around 10 glomeruli are necessary to achieve 90% performance (10.6 ± 0.5 mean ± s.e.m., *n* = 13 mice). Despite this smaller number of necessary glomeruli, they are not just a subset of the ones used in the fully saturated case.

Although our analysis showed that multiple glomeruli are necessary for detecting the target odors from glomerular inputs that incorporate our estimates of trial to trial variability and saturation, we asked whether single glomeruli may be sufficient to detect the targets making less conservative assumptions. We used receiver operating characteristic (ROC) analysis, a standard technique in signal detection, to estimate the ability to decode using single glomeruli, assuming linear summation of glomerular inputs and no trial to trial variability. We found that even in these favorable conditions, the best glomerulus-target odor pairing leads to a performance of less than 75% (with top 1-percentile 68.0%, and mean ± SD performance 52.8 ± 0.04%, *n* = 5473 glomerulus-odor targets pairs; Fig. 9C-D).

Since synaptic transmission is inherently noisy and plastic in the nervous system (del Castillo and Katz, 1954), an important question is how robust the decoder is to fluctuations in the readout weights. To test for robustness, we perturbed the classifier weights (for *C* = 10^6^) by multiplying them by a random factor that is normally distributed with a mean of one. We varied the perturbation level by varying the standard deviation of this factor. We found that the linear decoder is fairly robust to changes in its readout weights and that robustness is higher for the linear mixing case (*λ* = 1, Fig. 3D). For the data from all 13 mouse experiments, performance remained better than 90% of optimal at perturbation levels that are less than 0.25 for *λ* = 1, and 0.1 for *λ* = 0. However, robustness of classifier performance strongly depends on the specific target odors.

### Learnability and experience dependency of readout weights

We established that this task is linearly separable, but even a linear decoder has as many degrees of freedom as a mouse glomeruli. This raises the question how difficult it is to learn the readout weights; that is, how many examples are needed to properly constrain the weights; To quantify the need for learning, we first computed the odds of a ′random′ readout to perform well. Random readout weights were drawn from a Gaussian distribution with the same mean and standard deviation as the optimal linear readout weights. We evaluated the performance of 1,000,000 such readouts per target pair used in the experiment assuming *α* = 0.1 and saturating glomerular responses (*λ* = 0), always using the experimental trial structure (Fig. 4A). The average performance was 58.41 ± 4.53% (mean ± SD). The chance that a random readout performs better than 80% is around 1 in 10^5^. Thus, random hyperplanes typically fall short of separating the mixtures with target odors from the ones without. We next directly computed how many training samples are required to find the proper readout weights for performing the task. To find the minimum number of trials needed to generalize to the rest of the data, we varied the absolute number of training trials directly. With *α* = 0.1, only ~ 30 random trials were needed to get to 90% of asymptotic performance assuming linear mixing (*λ* = 1), yet ~ 150 trials for saturating mixture responses (*λ* = 0) (Fig. 4B). This difference highlights how saturation makes the task harder to learn, but crucially these numbers of training trials compare favorably to mice - typically mice had an order of magnitude more trials to reach sufficient performance before being tested with uniform distractor distributions (Rokni et al., 2014).

**FIG. 4.**
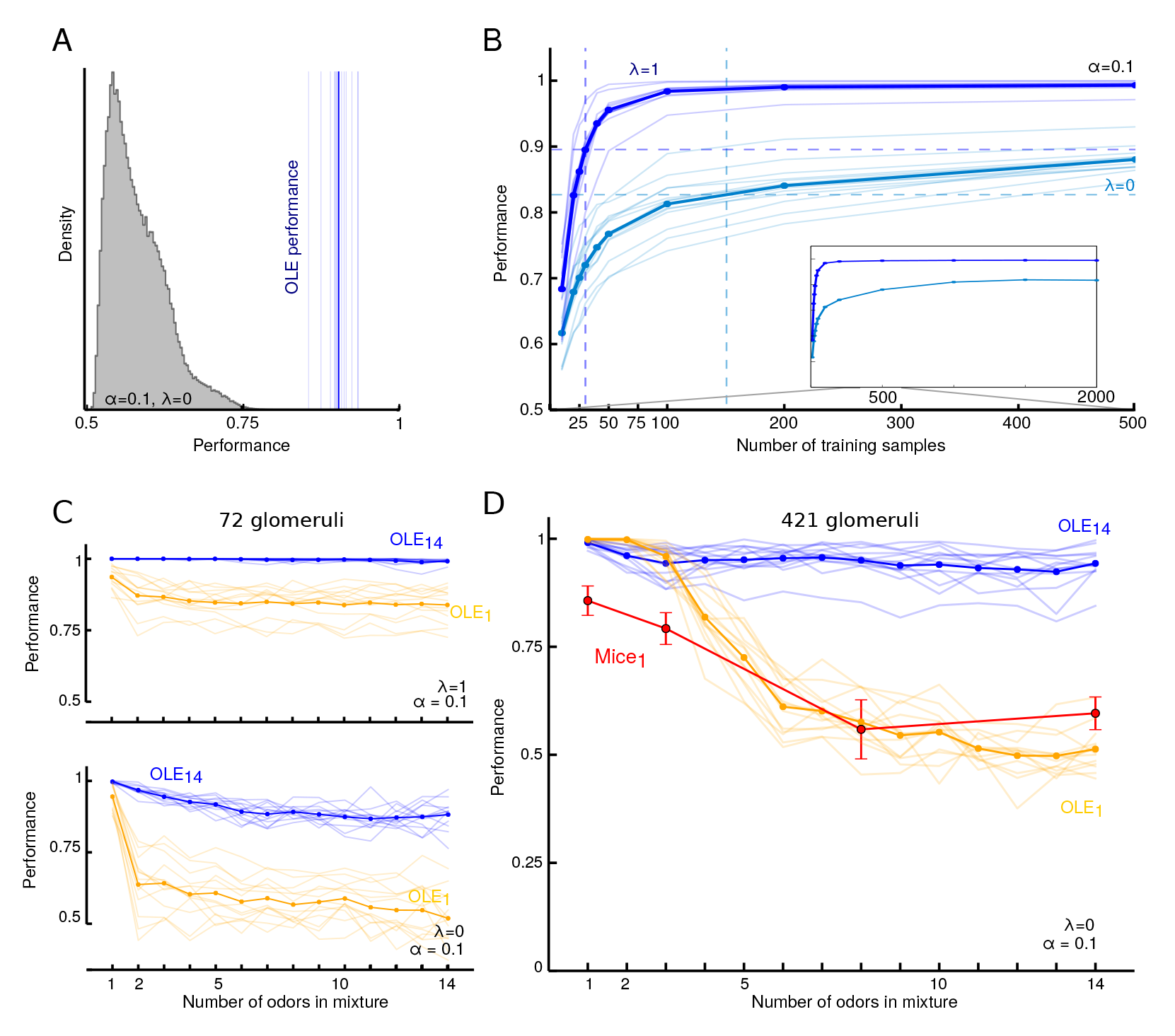
Learnability and dependence on trial statistics. (A) Distribution of performances for 13, 000, 000 random linear decoders with weights drawn from the distribution of OLE-weights and corresponding optimal threshold *θ*. The ability of 1, 000, 000 random readouts to detect the presence of the targets was evaluated for each of the 13 target odor pairs. The blue lines show the performance of the optimal linear decoder (OLE). (B) Performance vs absolute number of training samples for *α* = 0.1 and *C* = 106. For each target the average performance when trained on a small, random subset of samples is shown; each curve is averaged over 20 random training sets. The inset show the asymptotic average performance, which is reached when trained by 1, 500 - 2, 000 trials. The dashed, horizontal lines indicate 90% of the asymptotic performance, and the corresponding vertical lines mark the necessary training samples to achieve this performance. (C) Top: Average performance curves for OLE per target pair trained only on 80% of the single odor stimuli (yellow, OLE_1_), and on 80% of all the data, (blue, OLE_14_), with 0.1 noise and linear mixing based on 72 glomeruli. Bottom: same as top for non-linear mixing. (D) Average performance curves for OLE per target pair when trained only on 80% of the single odor stimuli (yellow, OLE_1_), and on 80% of all the data, (blue, OLE_14_) based on 421 glomeruli. The red curve shows the average performance ± s.e.m. of 5 mice trained on single odors and subsequently tested on mixtures of 1, 3, 8 and 14 odors (Mice_1_).

The analysis above indicates that a linear decoder can generalize from a rather small number of random mixture stimuli to thousands of other mixtures. Due to nonlinear mixing, this ability to generalize beyond trained samples strongly depends on the quality of training samples. A particularly meager training set is given by samples of just single odor components. Therefore, we trained classifiers only on single odors and tested them on arbitrary mixture stimuli. Without saturation the OLE performance for mixtures with more than one component degrades only mildly, but is substantially worse than an OLE trained with a training set containing all component numbers (Fig. 4C, top). With saturation, the performance of the OLE degrades strongly with increasing number of odors, and already for mixtures of two components has a performance of below70% (Fig. 4C, bottom). This inability to generalize is obviously stronger when fewer glomeruli are used for decoding. When we pool over all measured glomeruli (6 OBs, 421 potentially redundant, dorsal glomeruli), the performance of the OLE decays more gracefully with the number of components (Fig. 4D). Thus, for the linear readout, the mixture model, as well as the training stimuli, strongly affects the ability of the decoder to detect target odors.

### Discriminative learning vs. generative modeling

Complementary to the purely discriminative learning considered above, an important idea in neuroscience is that neural representations are also formed in a task-independent, unsupervised way by learning a generative model of the sensory input (Friston, 2010, Hinton, 2007a, Hinton, 2007b). The crucial advantage of generative modeling is the ability to learn tasks with much smaller number of task-specific training samples by exploiting general knowledge about the nature of the world. For our demixing task specifically this would mean that the glomerular representation (Fig. 1B) is first demixed into a representation akin to the one in Fig. 1A. Ideally this would allow the animal to perfectly learn the task from samples of just single odor components without the drop in performance observed for purely discriminative learning (cf. Fig. 4C-D).

To test the ability of mice to generalize beyond single odor components, we trained 5 mice for several sessions on the single odor task until they reached > 80% performance (see Methods). During test sessions, we mainly presented single odors, but in 10% of the trials, we also presented mixture stimuli with 3, 8 and 14 components (equally distributed). During these test sessions, mice continued to perform well for single odorant trials, and for (novel) mixtures of three. For mixtures with more components performance dropped substantially (Fig. 4D), consistent with OLE predictions. This pattern was observed in all mice individually (Fig. 10). Over the ~ 50 test stimuli per mouse and mixture number, their performance from the first ten to the last ten trials did not strongly fluctuate (2 significant increases, 1 significant decrease and 12 stable performances with 5% significance level; Mann-Whitney U test, p-values in Table I).

Our earlier experiments demonstrated that mice can learn this task when exposed to training samples with more complex odor components (Rokni et al., 2014). This suggests that due to the nonlinear nature of odor mixture encoding, both mice and OLEs have difficulty generalizing much beyond the trained complexity, and rather settle on decision boundaries that are good enough to solve the task. It further indicates that mice cannot significantly benefit from a generative model by which they could task-independently decompose odor mixtures into their individual odor components.

**TABLE I.** Comparison test trials of performance (p-values)

|  | mouse 1 | mouse 2 | mouse 3 | mouse 4 | mouse 5 |
| --- | --- | --- | --- | --- | --- |
| 3 odors | 0.028 | 1.000** | 0.851 | 0.504 | 1.000** |
| 8 odors | 0.502 | 0.495 | 0.927 | 0.912 | 0.966 |
| 14 odors | 0.500 | 0.931 | 0.500 | 0.922 | 0.009* |

P-values for Mann-Whitney U test with Monte Carlo permutation test statistics, i.e. fraction of cases where MWU statistic for shuffled data is larger than MWU statistic in last 10 and first 10 trials per mouse and odor condition. Labels: \*...significant performance decrease, \*\*...significant performance increase.

### Capacity of linear readout

Our analysis demonstrated that the presence of a single odor in mixtures of up to 14 odors is explicitly available for a linear decoder from the rich repertoire of receptors. What is the capacity limit for such a decoder? We can estimate this by using surrogate odor representations based on the representations of the 16 odors we measured. Surrogate odors were generated by drawing with replacement from the activations of the individual glomeruli by the 16 odors. We then calculated how well a linear decoder can report the presence of a pair of target odors in mixtures of up to 128 odors. Performance of classifiers was dependent on the target odor but all classifiers performed above chance level even for 128 odors, and above 80% for mixtures of around 32 odors (Fig. 5A). This is a remarkable ability since the classifier only uses 72 glomeruli out of the 3, 000 or more glomeruli in mice (Richard et al., 2010). We repeated the same analysis using pooled glomerular data from 6 olfactory bulbs (421 glomeruli). With this large number of (potentially redundant) inputs, the performance of classifiers was above 70% for mixtures of up to 128 odors (Fig. 5B). This analysis suggests that the capacity of mice to detect target odorants from mixtures may be well above what was tested.

## Discussion

An important task of the olfactory system is to identify odors of interest that are embedded in background mixtures. We formulated this task as a classification problem and asked whether a simple, biologically-plausible classifier can solve this task in the face of noise and non-linear integration of odorants at the level of olfactory receptors. To tackle this question we first determined an encoding model for mixture responses of olfactory sensory neurons as the input for the classifiers. The model is based on data from calcium imaging and generates mixture representations that are saturating sums of single odor representations with trial-to-trial variability. We find that olfactory receptors have a substantial capacity to transmit information about the composition of odor mixtures despite saturation and noise, which is explicitly available for a linear, multiglomeru-lar readout.

**FIG. 5.**
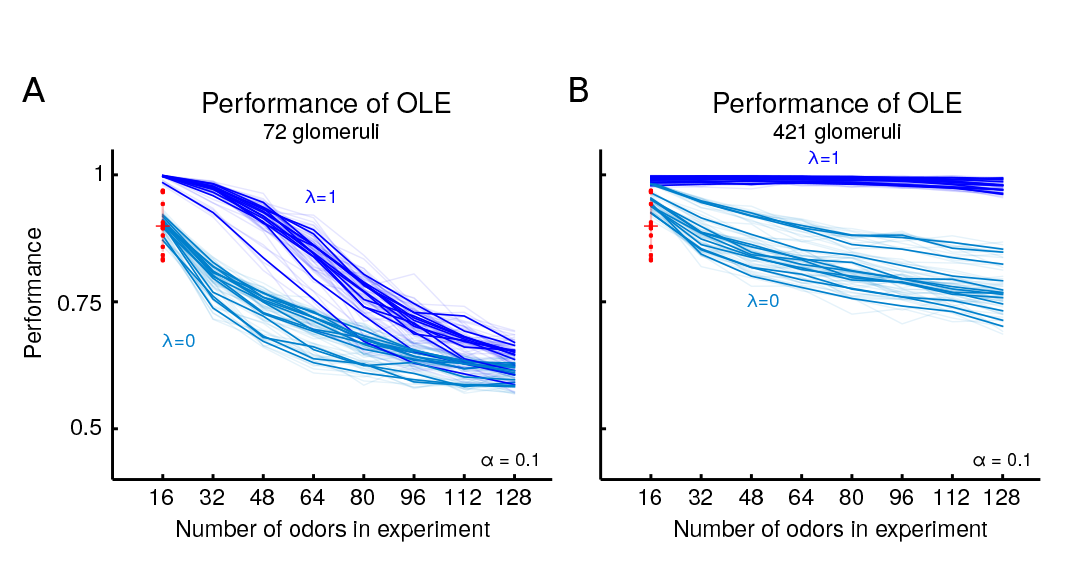
Capacity of an optimal linear decoder in cocktail party tasks with increasing odor numbers. (A) Performance of optimal linear decoder with linear and nonlinear mixing and noise *α* = 0.1 as a function of the number of odors. Additional odor patterns beyond 16 are generated by sampling from the responses of the 72 glomeruli to the 16 odors as described in the main text. Each faded line represents such a randomly generated odor data set and the 13 solid lines are the averages of 4 such lines with he same target odors. The red dots highlight the experimental performance of mice (B) Same as in panel A, but calculated by pooling across imaging experiments to form odor patterns with 421 glomeruli.

In recent years, object recognition has been identified as a core challenge to understand the visual system (DiCarlo et al., 2012). For (visual) object recognition, simple linear readouts are also sufficient to match behavioral performance (DiCarlo et al., 2012, Majaj et al., 2015) However, these linear readouts cannot be applied directly to photoreceptors, but rather to the neuronal activity of higher visual areas, after several nonlinear transformation steps, which are generated from multiple feedforward steps of processing (DiCarlo et al., 2012). Our finding that the identity of odors in mixtures can be directly decoded by linear readouts from glomeruli highlights a fundamental contrast between the olfactory and visual systems which is also mirrored anatomically: olfaction has substantially fewer anatomical stages than vision (Gire et al., 2013).

### Mixture Model

The general question we have posed in this study is whether glomerular responses to mixtures that include target odors and those that exclude target odors are separable. Due to the large number of possible stimuli in the behavioral task (> 10,000 possible mixtures), we estimated a statistical model of mixture response based on measurements made from a sample of mixtures. We found that the relationship between glomerular responses to mixtures and the responses to mixture components can be described by a linear sum followed by a saturating nonlinearity. For the odor stimuli we used in this study, suppressive or supralinear responses were not significantly above response variability (Duchamp-Viret et al., 2003). We also systematically measured the trial-to-trial variability in glomerular responses. We found two forms of response variability: one that is correlated across glomeruli and can probably be attributed to variation in the level of anesthesia and breathing, and another that was uncorrelated across glomeruli and may represent trial-to-trial variability in individual input channels. The magnitude of the uncorrelated noise was, on average, 10% of response amplitude. In anesthetized rats, the CV of single OSN firing was found to be on the order of 1 (Duchamp-Viret et al., 2005). Single glomeruli receive a few thousand sensory axons although this number can vary significantly depending on receptor gene (Bressel et al., 2016). If OSNs were statistically independent of each other, the glomerular CV is expected to be on the order of 1 - 2%. The higher estimate in our study may indicate that sensory neuron responses co-vary and are not truly independent, but may also reflect non-biological sources of variability in our measurements.

### Single glomeruli are not enough

In some cases, individual odorant receptors are thought to directly signal the presence of individual odors in a ′′labeled line′′ manner (Dewan et al., 2013, Ferrero et al., 2011, Hussain et al., 2013). These are thought to be involved in detecting specific odors that are important for innate behavior (Kobayakawa et al., 2007, Liberles, 2015, Stowers et al., 2013). Here we find that individual glomeruli are not sufficient to report the presence or absence of individual odors. Although it is possible that our limited sample of dorsal glomeruli miss more specialist glomeruli with highly selective responses, it seems implausible that there will be such glomeruli for a vast array of odors. Indeed, experimental evidence strongly supports the idea that most odorant receptors in the main olfactory system are broadly tuned (Godfrey et al., 2004, Young and Trask, 2002, Zhang and Firestein, 2002). Although individual glomeruli do not carry enough information to separate stimuli that contain a target odor from those that do not, we could reliably make this distinction by integrating information across different glomerular channels.

### No preprocessing required, a linear readout is sufficient

Our decoders were based directly on the nonlinear, noisy glomerular inputs. Other than removing cofluctuations across glomeruli, no intermediate stages of preprocessing such as normalization or decorrelation were necessary. Our analysis suggests that correlated trial-to-trial variability probably reflect slow changes due to anesthesia (Ecker et al., 2014) and breathing. Breathing induced covariation could be easily removed by lateral interactions (or to some extend by adjusting linear readout weights), which is why we neglected them here (Friedrich and Laurent, 2001, Olsen et al., 2010, Zhu et al., 2013, Blauvelt et al., 2013). Our measurements as well as those of others (Duchamp-Viret et al., 2005) indicate that noise levels at the glomeruli and overlap of glomerular patterns are not high enough to require an additional processing stage. The optimal linear readout relies on weights that reflect the pattern of glomerular activation by target odors and are orthogonal to the pattern for distractor odors. This mechanism inherently removes overlap of glomerular patterns. Normalization and decorrelation may become more important for large amplitude stimuli that may saturate neuronal elements more strongly or for suboptimal readouts (like template matching). Decorrelation of different types has been proposed in the olfactory bulb, most notably in the context of reducing overlap among patterns of activity for different odors (Friedrich, 2006, Gschwend et al., 2015). Such pattern decorrelation or separation is thought to help downstream readout when noise limits separability (Friedrich, 2006, Wilson, 2009). In the future it will be interesting to look at the role and amount of noise correlations in glomeruli of awake animals performing this figure-ground segregation task.

### Weights can be sparse but not random

Although our intent was not to make specific analogies with neural circuits in the mouse brain, it is tempting to compare the decoder readout weights to the connections from the OB to the piriform cortex (PC). Since PC neurons receive a sample (possibly random) of glomerular input (Ghosh et al., 2011, Miyamichi et al., 2011, Sosulski et al., 2011), we also asked whether imposing sparseness constraints on the decoder affects performance. Remarkably, less than 20 glomeruli were sufficient to achieve performance that matches mice. It is likely that even fewer glomeruli may suffice if we were able to choose from the entire complement of mouse glomeruli. We found that while sparse connectivity was sufficient for robust performance after adequate training, classifiers built with random connectivity based on the same statistics as the optimal linear readouts were typically not successful. The problem remains linearly separable within the higher dimensional PC representation, but crucially this recoding step is not necessary for a linear readout to be successful. Furthermore, our analysis of random readout suggests that even in a large PC population there might only be a few neurons that are selectively tuned to the target odors, due to the large number of mixtures. To the extent we can make an analogy to mouse olfactory circuits, our findings suggest that any stochastic connectivity from the OB to PC may have to be supplemented with synaptic modification in the inputs, associative fibers or outputs of the PC to allow odor mixture analysis. Whether such learning occurs associatively over an extended period as animals experience odor environments, or whether it requires more specific reinforcement is an interesting question for future experiments. Alternatively, learning could happen via granule cells that dynamically gate the appropriate glomerular channels via mitral cells to the PC (Koulakov and Rinberg, 2011, Markopoulos et al., 2012), which would predict the existence of target-odor specific feedback modulation of mitral cells.

Although the optimal linear readouts are sufficient to match the performance of mice given the encoding model, we also showed that SVMs, which can approximate arbitrarily complex decision boundaries, outperform those simple decoders and suffer from decaying performance for increasing numbers of distractors. This performance decay suggests that for both mice and machines the ultimate bottleneck in this task is the overlapping glomerular representation. While SVMs were not required to mimic mouse performance, it is conceivable that the myriad parallel pathways from the OB could implement decoding schemes similar to an SVM and that over time plasticity rules allow for the learning of intricate decision boundaries; however, for this segregation task nonlinear decision boundaries are not necessary.

### Robustness and training dependence

Other than the overall performance, the difficulty of classification can also be assessed by the robustness of classification to modifications in readout weights and by the speed at which a classifier converges on the proper weights. Even linear classifiers can learn the task relatively easily in dozens of trials, despite response variability and saturation. Although a direct comparison of learning rates of classifiers and mice may not be appropriate, it is worth noting that mice take hundreds of trials to learn the task under our conditions. We also find that once a classifier has been trained, small perturbations of the weights did not strongly affect performance. This indicates that the classifier has robust performance in a local region of the weight space, which nevertheless cannot be readily reached by random assignment. Our analysis of random readouts indicates that there is a 1 in 10^5^ chance of having a performance of above 75%. Extrapolating this insight to neural architecture of the mouse olfactory system, it might mean that any projections that start out randomly (e.g., the OB to cortex projections) will need to be modified with learning, but that there might be several ′template′ neurons whose high correlation with rewards will facilitate learning.

We showed that optimal linear readouts based on glomerular activity are highly sensitive to the quality of the training set. When trained on single odors, they perform poorly for mixtures. An ideal observer that knows how individual odors mix should perform accurately for mixtures even when trained on single odors. To be specific, assuming an explicit δatomicδ representation of the single odors in the olfactory cortex, and that a decoder has access to this population, the readout should generalize well to mixtures when solely trained on the atoms. This reflects the main advantage of generative modeling, which has therefore been hypothesized to play an important role for shaping sensory representations (Barlow, 1997, Friston, 2010, Hinton, 2007a, Hinton, 2007b) We tested this prediction in mice and found that while their performance for single odors remains stable and their performance for mixtures of three is comparable, mice generalize poorly to mixtures of eight and fourteen. This suggests that mice learn the segregation task by adaptively adjusting decision boundaries directly based on incoming sensory information rather than highly processed, demixed representations. Due to the highly nonlinear representation of mixtures, such decision boundaries will generalize poorly when only trained with single odors. An alternative explanation for the mouse behavior (i.e., failure to detect target odor in mixtures when trained only on single odorants) is that they learned a different task rule - for instance, that two of the 16 odors are rewarded only when presented alone. Due to the novelty and low relative frequency of the mixture stimuli they may have ignored those stimuli and thus failed to generalize. Such an interpretation however is at odds with the good performance for mixtures of three odors and the fact that their errors consisted of both false alarms and misses. The training data from our previous study also argue against this alternate interpretation (Rokni et al., 2014). The mice in that work were trained by picking odor mixtures with increasing complexities - from distributions ranging from few distractors to uniform distractor distributions. Whenever a mouse reached 80% performance, the distribution was changed (in three steps until the uniform distribution was presented in the test phase). There the mice required > 1,000 trials to reach this level, and usually also encountered hundreds of trials per condition. This experience dependency dwarfs the small number of trials that the linear decoder needs to learn the task. However, carefully designed future experiments are necessary to address the question of how much training is required for learning in mice. Arguably, learning the task relies at least on two independent components. First, mice have to learn the task structure (go/no go, rewards, timing, etc.) and second mice have to learn which target odors are rewarded. By using the same task structure for different odor sets, one could greatly reduce the learning phase for the structure part and better estimate the number of trials that mice need for learning the task.

Experiments similar to those in the generalization task we studied in this paper (Fig. 4D) have been done in humans. Jinks and Liang trained humans initially for 3 days on single chemicals and then tested them for mixtures (Jinks and Laing, 1999). The subjects detected single chemicals with performance of around 70% and their performance for mixtures of 12 dropped to chance level. Our modeling, as well as our own experiments, suggest that this decay in performance is both a consequence of the training data and the synthetic encoding of stimuli, which makes generalization difficult. In vision, the intriguing characteristics of training schedules has long been noted - i.e. a minimal number of training examples is needed per session for learning and an interleaved stimulus presentation design can hinder learning (Aberg and Herzog, 2012; see also ; see also Lechner et al., 1999 for nonvision examples). Consequently, we believe that the study of the impact of training schedules could also be fruitful for understanding olfactory circuits in mice.

### Capacity

Overlapping representations of odors allow the olfactory system to encode more odors than the number of receptor types (Hopfield, 1999). However, broad tuning of receptors may decrease the discriminability of stimuli when noise and saturating glomeruli are considered. An earlier analysis of the coding capacity of spatial patterns of glomerular activation (Koulakov et al., 2007) pointed out that even with all-or-nothing glomerular activity, it is possible to simultaneously represent a dozen mixed odorants. Our analysis considers graded glomerular responses and empirical measurements of responses to specific odors and suggests that the capacity of the system is substantially greater. Whether this estimate is comparable to the capacity of mice remains unknown. Yet, understanding the capacity to detect and discriminate odors (Bushdid et al., 2014, Meister, 2015, Weiss et al., 2012) will provide crucial insights for understanding the olfactory system.

### Conclusions

In this work, we have linked experimentally-measured glomerular responses to behavioral data in an odor demixing task. Using realistic assumptions about neuronal noise, and nonlinear interactions for mixtures, we show that the information about odor-mixture components at the level of olfactory receptors is already linearly separable and does not require any preprocessing or inference algorithms that rely on prior information and feedback circuits.

## METHODS

Computational analyses are based on published behavioral and imaging data (Rokni et al., 2014). Further imaging experiments to estimate trial-to-trial variability and mixing properties in glomeruli as well as a behavioral test for generalization have been performed as described below. All procedures were performed using approved protocols in accordance with institutional (Harvard University Institutional Animal Care and Use Committee) and national guidelines.

### Summary of behavioral task and data

An odor panel of 16 single monomolecular odors was used and for each mouse two odors were selected as target odor pair. In each trial, one of these target odors was presented with 50% probability, together with a random selection of distractor odor
s, whose number is uniformly distributed. Mice were trained to report the presence of the target by licking a water spout and its absence by withholding licking. Licking on go trials (hit) was rewarded with water, whereas licking on no-go trials (false alarm), as well as withholding on go trials, was punished with a time-out which increased the time to the next trial. Mice learned to perform this detection task with above 80% performance after being trained over just a few hundred trials, with some mice performing close to 100% (Rokni et al., 2014). Each mouse was trained on a fixed pair of target odors and after training tested over multiple days with the mixture statistics described above. For each mouse, we selected the behavioral trials on each day starting from the first lick and ending with the last lick (which resulted in dropping about 10 trials per session) and concatenated those trials to create a continuous behavioral set, describing which odor was present and what behavior the mouse performed. There were 4 - 10 such sessions per mouse, which lead to behavioral data sets with 1, 500 - 3,500 trials each, for 13 mice. Some of these mice had the same target odor pairs.

### Novel single odor task

The procedures for behavioral training were performed similarly to the ones described elsewhere (Rokni et al., 2014). In brief, 5 c57bl6 mice (Charles River) were first anesthetized (Ketamine/Xylazine 100 and 10 mg/kg, respectively) and a metal plate was attached to their skull with dental acrylic for subsequent head restraining. Following 1 week of recovery from surgery, mice were water deprived and were trained on the task. During training a single odorant was presented for 2 seconds every 10 seconds and mice had to respond correctly within the 2 second period. Licking when a target odor was presented was rewarded with a 10*μ*l water drop, correct rejections were not rewarded, and incorrect trials were punished by a 5 second timeout. Half of the trials were go trials in which one of the two target odors is presented, and half were no-go trials in which one of 14 background odor is presented. Testing sessions began after mice performed above 80% correct for a whole session. In testing sessions, a mixture of 3, 8, or 14 components was presented every 10th trial with a 50% probability of a target odor being present. Mixtures with different numbers of components were equally presented. Once the number of components in the mixture was determined as well as the trial being a go or a no-go trial, the specific composition of target and background odors was random. The individual test performance curves are shown in Fig. 10.

**FIG. 6.**
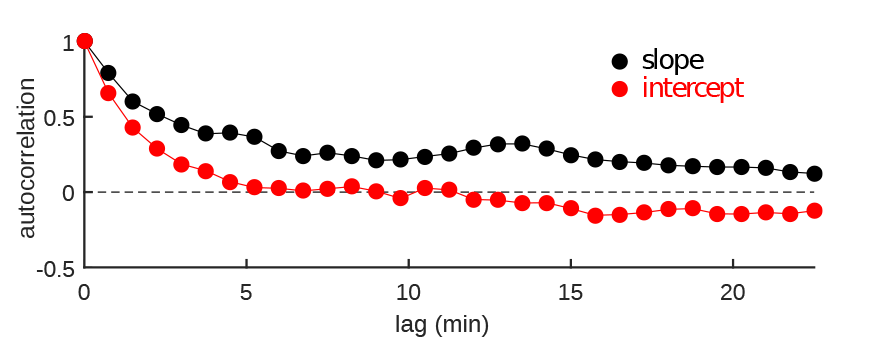
Timescale of changes in correlated glomerular response variability A linear fit was used to describe the relationship between individual responses to each single odor presentation and the mean response to that odor (Fig. 1G). To measure the timescale at which the parameters of the linear fit change, we plot the autocorrelation of the slope (black) and intercept (red) of these fits. The slow changes are in line with correlated variability being related to anesthesia.

### Glomerular Imaging

Adult OMP-GCaMP3 mice (Isogai et al., 2011) were anesthetized with ketamine/xylazine (100 and 10 mg/kg, respectively) and the cranial bone over the OB was removed with a dental drill. The exposed brain was covered with a thin layer of isotonic agar and a coverslip. An Olympus objective (10X, NA 0.3) was used to image the OB surface onto the sensor of a CMOS camera (DMK 23U274, The Imaging Source GmbH). Images (1600 × 1200 pixels, approximately 750 by 560 microns) were acquired at 8 bit resolution and 7.5 frames/sec. Data from the camera was recorded to the computer via data acquisition hardware (National instruments) using a custom software written in LabView. A blue LED (M470L3, Thorlabs) was used for excitation. Excitation and emission light were filtered using a filter set (MDF-GFP, Thorlabs). Odors were delivered using an automated olfactometer that is designed to have constant flow and to have each odor independent of all other odors (Rokni et al., 2014). All odorants were diluted to 30% v/v in diethyl phthalate and then further diluted 16 fold in air. The 16 odors and their mixtures were presented in random order. Odors were presented every 45seconds for 2 seconds with data acquisition beginning 3 seconds before and ending 3 to 5 seconds after odor presentation.

### Glomerular image analysis

Glomeruli were identified using two different methods. First, the response to each individual odor was converted into a *dF/F* image (using averages of all frames during odor presentation and average of all frames in baseline period) and *dF/F* images in response to repetitions of the same odor were averaged. The 16 mean *dF/F* images were then collapsed into a single image by taking the maximal intensity projection. Putative glomeruli were then marked by hand for further analysis. In a second method, the response in each trial was converted to a *dF/F* image. Each odor response was then represented as a point in pixel space and principal component analysis was used to find covarying pixels. Putative glomeruli were hand drawn based on the first 8 principal component images. For each experiment, we chose the method that yielded more putative glomeruli.

Once putative glomeruli were selected, the mean value in each frame was calculated to obtain time varying response traces that were then converted to *dF/F* traces. Response magnitude was quantified by integrating these *dF/F* traces. Variability of single odor responses was measured as the coefficient of variation (CV) of glomerular responses, also denoted by *α*. We found that responses had two sources of variability - a common variability that affect all glomeruli, and private variability for each glomerulus across trials. We reasoned that the common variability was likely linked to global changes relating to fluctuations in anesthesia and breathing, and chose to remove it. We first built a vector of responses of all individual glomeruli for each odor. For each of the 16 odors, we calculated the best linear fit between the vector for each trial 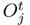 and the average vector for that odor *O*_*j*_. We then subtracted the best fit, and calculated the remaining (private) variability for each glomerulus, i.e.

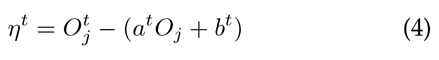

Where *η*^*t*^ is the deviation of glomerular response from the linear fit, 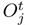 is the response to the *j*th odor on trial *t*, *O*_*j*_ is the trial-averaged response to odor *j*, and *a*^*t*^ and *b*^*t*^ are the slope and intercept of the best linear fit between 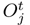 and *O*_*j*_. For each glomerulus *i* we then calculated the standard deviation of *η*_*i*_, and plotted it against its mean response amplitude *O*_*ij*_. The non-correlated CV for each experiment was measured as the slope of the linear relationship between the non-correlated SD and response amplitude (Fig. 1H).

Summation of mixture components was analyzed by comparing mixture responses to the linear sum of the mean responses to mixture components. The data were fitted by a sigmoidal function (for each glomerulus and mixture):

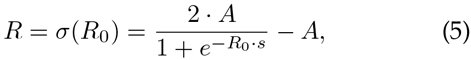

where *R* is the response to a mixture, *A* is the saturation level for the glomerulus, *R*_0_ is the linear sum of mixture component responses, and *s* is a free parameter.

### Mixture models for decoding analysis

For the decoding analysis, we considered the following encoding model, which is based on our measurements. Given the average glomerular response vectors *O*_*j*_ and the trial structure *c*_*j*_(*t*), i.e. a binary variable denoting the presence of odor *j*in trial *t*, the mean (linear) glomerular response pattern to a mixture in trial *t* is given by:

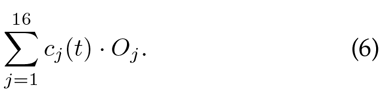

This mean response is subject to trial-to-trial variabity. The linear response *R*_0_ is thus the sum of the s gle component response *O*_*j*_ with an added noise te *η*_*j*_ that is normally distributed with a mean of zero and a variance of 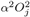 as defined in Eqn. (2). The factor *α* controls the coefficient of variation. The response *R* a mixture in trial *t* is a sigmoid function of the linea mixed trial response *R*_0_(*t*):

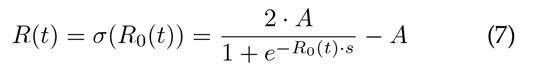

where *A* is the saturated response amplitude and *s* is a free parameter that governs the slope of this sigmoidal function. For each glomerulus *i* we assumed *A* to be equal to the maximum of the linear sum of component response 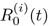 and set *s* = 10/*A*. This makes 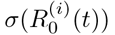 almost fully saturated for inputs larger than *A*/2 and the output level is around 50% of the saturation level for input values around *A*/10. To assess the effect of this nonlinearity, we tested encoding models that range from fully linear (*λ* = 1) to fully nonlinear (*λ* = 0) as defined in Eqn. (1).

### Decoding analysis

The glomerular responses in a given trial *t* are drawn from the probability distribution for response vectors *R*(*t*), which are generated using the mixture models discussed above. For any given trial *t*, the target was either present or not and we denote this ideal behavior by the binary variable *r*(*t*). We are interested in the performance of ideal observers that use different algorithms, like linear readouts and support vector machines, as specified below. This is a typical example of the problem of pattern recognition in statistical learning theory and can be formulated as a risk minimization problem (Hastie et al., 2009, Vapnik, 1999). Using the Kronecker-*δ* loss function *δ*(*r*, *f*(*R*, *θ*)) between behavioral response r and a given sensory input R, and the response of the readout *f*(*R*, *θ*) with parameters *θ* one can write the empirical risk functional for a subset of trials I with cardinality |*I*|:

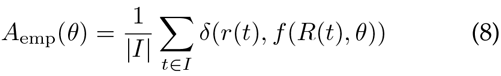

**FIG. 7.**
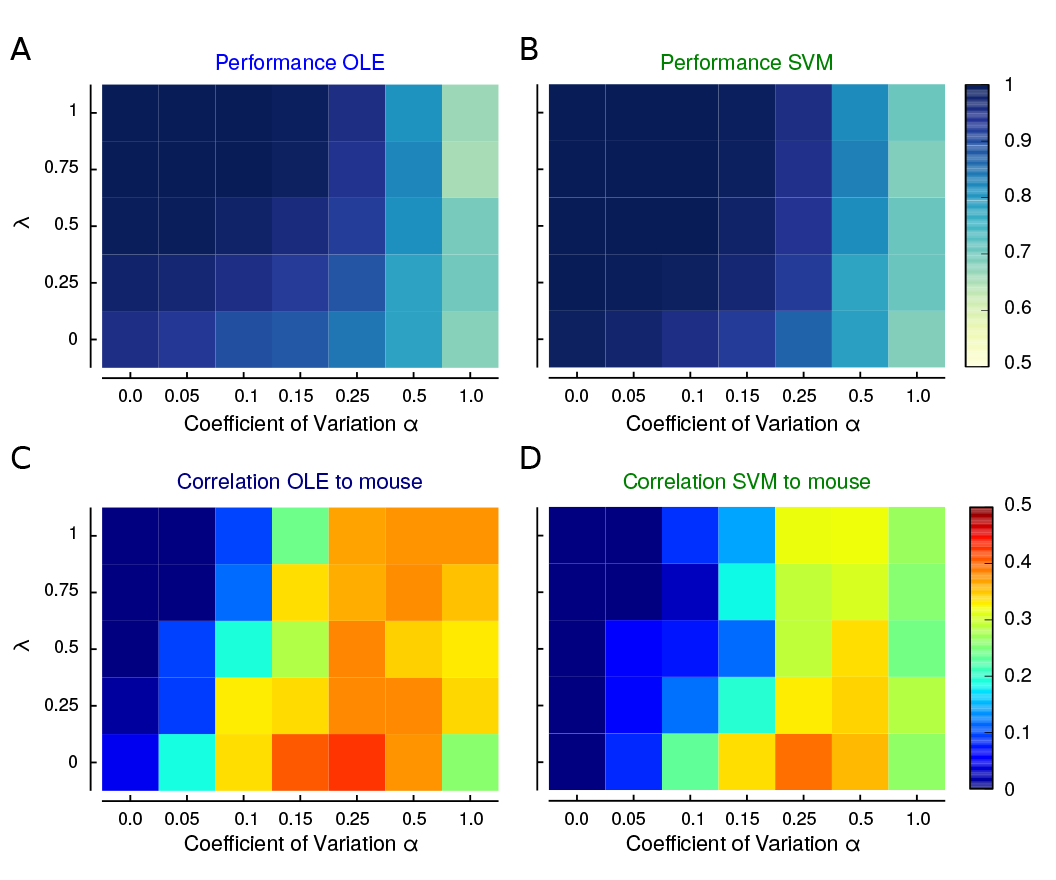
Comparing OLE, SVM and mice. (A) Performance of optimal linear decoder as a function of noise level and nonlinearity for one target odor pair hexyl tiglate and methyl tiglate. Average over all animals is shown in Fig. 2C. (B) Performance for SVM with radial basis function for mixture data like in (A). (C) Average correlation across mice, between performance curves of mice and OLEs for simulated pairs of parameters *λ* and *α*. The performance curves are best correlated for *λ* = 0 and *α* = 0025. (D) Same as (C) but for performance curves of SVMs. Highest correlation was also observed for *λ* = 0 and *α* = 0.25. These performance curves are shown in the main text (Fig. 2E-F).

Since the Kronecker-*δ* is one when both entries are the same, and zero otherwise this empirical risk function measures the fraction of correct classifications by readout *f*(*R*, *θ*). The goal is then to find parameters *θ*_0_ for the readout *f*(*R*, *θ*_0_) that minimizes the empirical risk on the training set I. To compare different readouts we compare their performance on the test set, i.e. the other trials that have not been used for training the readout. All read-outs are cross-validated using the repeated random subsample technique; unless otherwise specified we used 80% training data, 20% test data and 20 repetitions with randomly sampled training and test data sets. We report the average test performance, defined as the mean number of correct classifications of *r*(*t*) by *f*(*R*(*t*), *θ*_0_) averaged over the test set.

Specifically, we want to find the ′simplest′ algorithm (i.e., the readout function *f*) for the demixing task that matches the error of animals. As we describe below, we consider: algorithms based on single glomeruli, the optimal linear decoder, support vector machines (SVM) with radial basis functions and logistic regression.

**FIG. 8.**
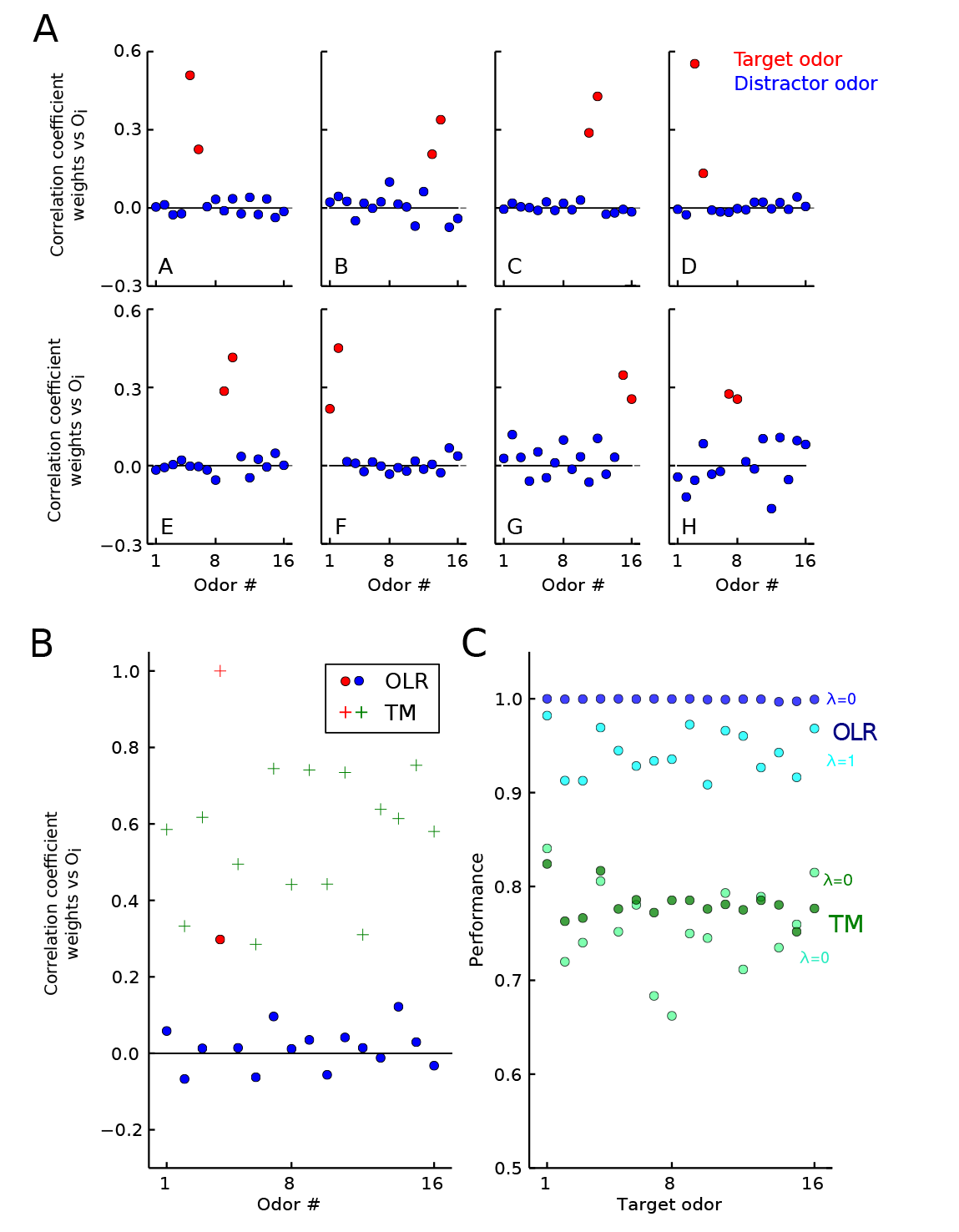
Structure of readout weights and template matching performance. (A) Correlation coefficients between the glomerular readout weights (for *M* = 10^6^, reported in Fig. 3C) and the 16 glomerular odor patterns for different odor target pairs (A-H). Some target odor pairs in Fig. 3C occur more than once, here only one example per odor pair is shown. The red dots highlight correlations to target odors and the blue dots to distractor odors. Generally, readout weights are highly decorrelated with distractor odor patterns and positively correlated to target odor patterns. (B) We illustrate the inadequacy of a simple template matching algorithm. This is best done with just one target odor, which we varied. For a given target odor the template matching algorithm (TM) uses the glomerulus pattern of this odor as a readout weight. Example correlation coefficients of the weights for target odor 4 to all odor patterns for both the OLR and TM are shown. As for target odor pairs, the OLR′s readout weights are uncorrelated to distractor odors. In contrast, due to overlapping odor tuning, the TM-readout weight is correlated to distractor odor patterns. Consequently, the TM-performance also suffers as shown in (C). (C) Performance for OLR and TM vs. target odor for linear (*λ* = 1) and nonlinear case (*λ* = 0) with *α* = 0.1. For the TM algorithm the threshold (*θ* in Eq. (3)) was optimized for performance. Nevertheless, the TM-performance is around 80%, significantly below the OLR′s performance.

### Single glomerulus decoder

When considering single glomeruli (Fig. 9), we calculated their performance without cross validation, which will overestimate their performance, and therefore provides an upper bound on their performance. Specifically, for each glomerulus *i* and pair of target odors we calculated their empirical histograms: *P*(*R*_*i*_(*t*)|*r*(*t*) = 0) and *P*(*R*_*i*_(*t*)|*r*(*t*) = 1). Based on these histograms we calculated the ROC curve and therefore the hit rate (HR) and correct rejection rates (CR) for varying thresholds *θ* (Green and Swets, 1966). We then reported the maximum performance for the best threshold.

### Linear decoder

The linear decoder depends on the following parameters: the readout weights *w* and the decision threshold, and has output *y* = *H*(*w*^*T*^ · *R*(*t*) + *θ*) (Eqn. (3)). The optimal linear weights (OLE weights), are the ones that maximize the empirical risk function, which is equivalent to minimizing the norm

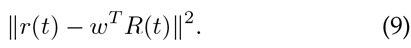

For the linear mixture models without noise, the weights that minimize this expression have a particularly simple form, as we will show first. In general, the optimal parameters for linear regression are given by the normal equations (Hastie et al., 2009): *R*^*T*^(*r* – *R* · *w*_OLE_) = 0, with *R*^*T*^ = (*R*_1_, *R*_2_, …, *R*_*N*_) and *r*^*T*^ = (*r*_1_, *r*_2_, …, *r*_*N*_) for all *N* concatenated trials. If *R*^*T*^*R* is not singular, the solution to this normal equation is given by:

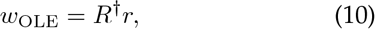

where † is the Moore-Penrose pseudoinverse. This formula holds irrespective of the mixture model. For the linear mixture model the matrix *R* is given by *R* = *MO*, where *O* is the matrix of odor patterns and *M* is the mixture matrix. Therefore, we get *w*_OLE_ = *O*^†^*M*^†^*r*. Due to the task structure, the rewarded trials are the sum of either of the target odors′ characteristic function, i.e. 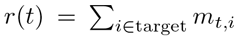. Thus, it follows that the *i*th entry of *M*^†^*r* equals 1 if *i* is one of the targets and 0 otherwise. Therefore, we obtain that 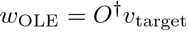, where *v*_target_ is a vector that is 1, when the *i*th index is a target and 0 otherwise.

**FIG. 9.**
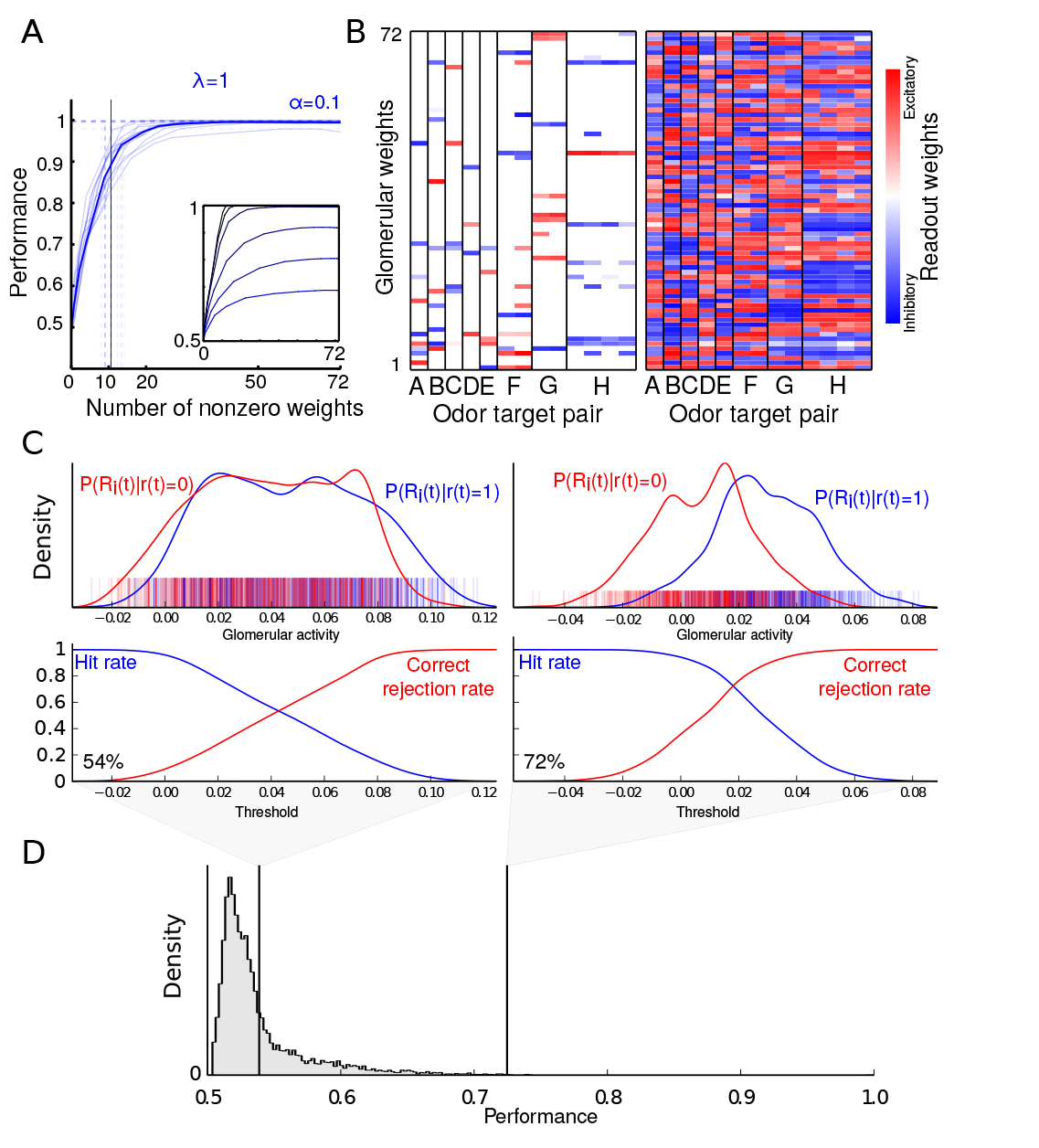
Decoding performance of single glomeruli and OLR without saturation. (A) Performance versus number of nonzero weights of the OLR (*λ* = 0). Each faded line depicts an animal and solid line depicts the average. The horizontal, dashed lines indicate asymptotic performance and the vertical dashed lines indicate the corresponding minimal number of glomeruli to achieve 90% of that performance per animal. Their average is highlighted as solid vertical line. Inset: Average performance vs. number of non-zero weights of the OLR for varying noise levels (*α* = 0, 0.05, 0.1, 0.25, 0.5, 0.75, 1, from top to bottom). (B) Example readout weights obtained for 90% of asymptotic performance (left) and asymptotic performance (right). Several of these 13 mice had the same target odor pairs, as indicated by grouping under the same letter. OLR learns similar readout weights from different trial statistics for the same target odors. Across different target odors, the readout weights vary substantially - but the weights are similar for the same target odors. (C) Top panels: Conditional distributions of glomerulus activity for the presence *r*(*t*) = 1 or absence of the target odors *r*(*t*) = 0 in the linearly mixed activity of two example glomeruli. The activity for each glomerulus is calculated as the linear sum of the average *dF/F* for individual components of that glomerulus. We estimated the probability density from the individual samples (shown as vertical lines) by kernel density estimation with a Gaussian kernel and width given by Scott′s rule. Bottom panels: Corresponding hit rate and false alarm rate curves for varying thresholds. The number indicates the performance for the best threshold. (D) Histogram of (best) performances for all glomeruli across imaging data sets and for all target odor pairs. The performances for the two examples from (C) are indicated by vertical lines. Most glomeruli have less than 70% performance.

All reported performances are the averages calculated by cross validation, which was used to determine the optimal threshold and weights. For each resampling, we calculated the OLE weights as described above and computed the optimal decision threshold *θ* as the one that maximizes the training performance on 80% of the data. This threshold is then used to predict the output for the test set (the remaining 20%), which is used to calculate the test performance. This procedure is carried out for 20 random resampling of the data and the average test performance is reported.

### Support vector machine

We studied the performance of support vector machines (SVM) with radial basis functions as kernels (Muller et al., 2001, Vapnik, 1999). As parameters *θ* for the SVM we performed a grid search over the penalty parameter for the error term *C* ∈ {10^−2^, 10^−1^, 1, 10, 10^2^, 10^3^, 10^4^, 10^5^, 10^6^} and the kernel coefficient γ ∈ {0, 1, 2, 5, 10, 20, 50}. The optimal parameters were determined on the training set and we report the average performance for these parameters on the test set.

**FIG. 10.**
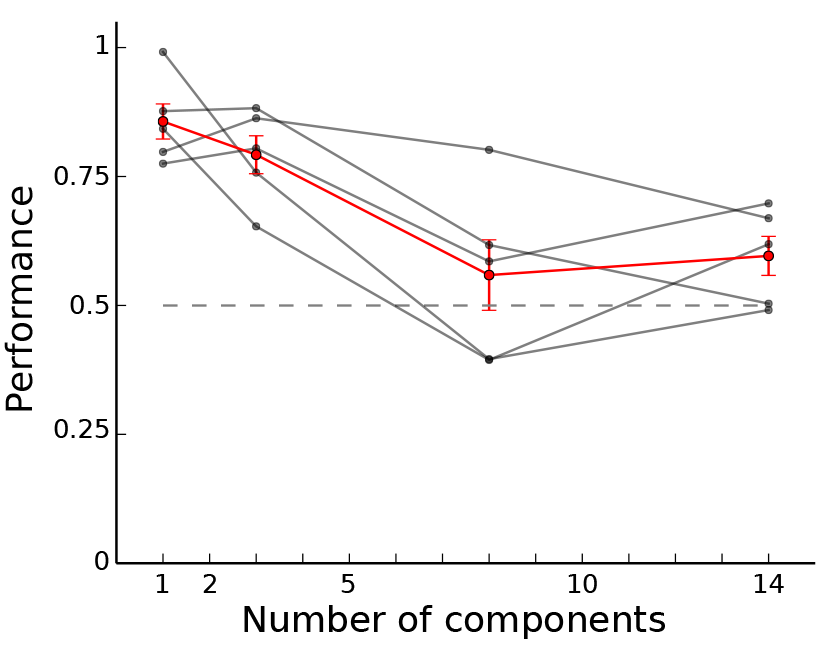
Individual performance curves. The red curve shows the average performance ± s.e.m. of 5 mice trained on single odors and subsequently tested on mixtures of 1, 3, 8 and 14 odors, and the gray curves show the performance curves for the 5 mice individually.

### Logistic regression

We used regularized logistic regression to decode the presence of a target odor. For binary classification the likelihood for the logistic regression is given by (Bishop, 2006):

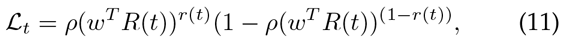

with logistic function *ρ*(*s*) = 1/(1 + exp(−*s*)). Minimizing misclassification is equivalent to minimizing the logarithm of the likelihood. Therefore, the error functional for the regularized logistic regression is given by:

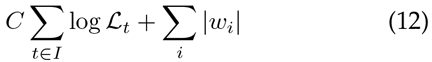

We use the L1-norm to bias towards sparse read-outs, and the constant *C* varies the degree of sparseness as it balances the trade-off between classification errors and the summed absolute readout weights. We varied the regularization constant from 10^−2^ to 10^6^.

We employed the Scikit-learn toolbox in Python to perform the SVM and logistic regression analysis (Pedregosa et al., 2011).

### Weight perturbation analysis

Initially, the OLR is trained on 80% of the data, to obtain the optimal weights *w*_OLR_ and the performance on the remaining 20% is reported for weight perturbation level 0. Then, for different levels of the weight perturbation (*wp*), each component of *W*_OLR_ is multiplied by a value drawn from a Gaussian distribution with mean 1 and variance *wp*^2^. The performance of the OLR with this perturbed weight is then calculated and averaged over 20 randomizations per perturbation level. The results are shown in Fig. 3C.

## AUTHOR CONTRIBUTIONS

A.M., D.R., M.B. and V.N.M. designed research; A.M. performed simulations and model analysis with input from M.B.; D.R. performed imaging and behavioral experiments. V.K. contributed glomerular maps. A.M., D.R. and V.N.M wrote the paper with input from all authors.

## ACKNOWLEDGMENTS

We thank Philipp Berens, Mackenzie Amoroso and Alexandra Ding for helpful feedback. This work was supported by Harvard University, by DFG grant MA 6176/1-1 (AM), Marie Curie Fellowship PIOF-GA-2013-622943 (AM). MB has received financial support from the Bernstein Center for Computational Neuroscience (FKZ 01GQ1002) and the German Excellency Initiative through the Centre for Integrative Neuroscience Tubingen (EXC307). Research in VNM′s lab is supported by grants from the NIH (DC011291, DC014453). Computational resources were provided and maintained by Harvard FAS Research Computing.

